# Gene co-expression network reveals key hub genes associated with endometriosis using bulk RNA-seq

**DOI:** 10.1101/2025.08.10.669560

**Authors:** Nooshin Ghahramani, Ali Hashemi, Seçil Eroğlu, Elaheh Esmaeili Kordlar

**Affiliations:** Department of Animal Science, Division of Animal Breeding and Genetics, Faculty of Agriculture, Urmia University, Urmia, Iran; Department of Medical Biology, Faculty of Medicine, Gaziantep Islam Science and Technology University, Gaziantep, Turkey; Chemical Sciences, Faculty of Chemistry, University of Padova, Padova, Italy

**Keywords:** *EMs*, *DEGs*, Meta-analysis, Pathway enrichment, Co-expression network

## Abstract

Endometriosis *(EMs)* is a complex and prevalent gynecological disorder with a significant genetic component, posing a major clinical challenge in reproductive medicine due to multifactorial inheritance patterns and the involvement of gene–environment interactions in pathophysiology. However, despite extensive research, reliable diagnostic biomarkers for *EMs* have yet to be identified. We utilized bulk transcriptome sequencing data obtained from the Gene Expression Omnibus to identify hub genes involved in *EMs*. This study was conducted using a system biology analysis, incorporating differential gene expression, meta-analysis of transcriptomic data, functional enrichment analysis, construction of gene co-expression networks, and comprehensive topological analysis to identify key regulatory genes. Bulk *RNA*-seq analysis revealed significant differential gene expression between healthy and *EMs* groups. Overall, 603 and 443 meta-genes were discovered using the *Fisher* and *Invorm P-value* combination methods, respectively. A total of 427 meta-genes were subjected to functional enrichment analysis, which revealed significant enrichment in several *KEGG* pathways related to *EMs* including *“Adherens junction,” “p53 signaling pathway,” and “AMPK signaling pathway.”* Additionally, Gene Ontology analysis revealed key processes including “Regulation of Anatomical Structure Morphogenesis,” Acetylglucosaminyltransferase Activity” and “Positive Regulation of Intracellular Signal”. Co-expression network analysis identified the turquoise module as a critical functional module, within this significant module, the genes *IGFBP7, IGFBP3,* and *NKAP* were identified as *EMs* hub genes based on high connectivity and central roles in the network. The constructed protein–protein interaction network further highlighted *STAR, PLCD3, RPAP2, MSI2, MAS1, TBX1, LIPT1,* and *SVIL,* as key genes. These genes represented high centrality within the network, suggesting potential regulatory and functional significance in the molecular mechanisms underlying *EMs*. Notably, *miR-143-3p, miR-340-5p, miR-410-3p,* and *miR-302b-5p* were implicated in *EMs*-associated regulatory networks. This integrative approach significantly enhances our understanding of the molecular mechanisms underlying *EMs* and provides a robust foundation for the development of diagnostic biomarkers.

## Introduction

Endometriosis *(EMs)* is a painful condition in which tissue that is similar to the inner lining of the uterus grows outside the uterus. *EMs* often affect the ovaries, fallopian tubes and the tissue lining the pelvis (1). Those with the condition often suffer from dysmenorrhea, dyspareunia, infertility, and pelvic pain, which negatively impact patients’ quality of life (2). *EMs* carries an increased risk of developing ovarian cancer, particularly clear cell carcinomas and ovarian endometrioid carcinomas (1). *EMs* is a multifactorial disease with complex host-immune interactions which causes Recurrent Implantation Failure *(RIF)*, affecting the endometrial receptivity and the embryo implantation process, leading to infertility and pregnancy failure (3). The immune system plays an important role in the beginning and progression of *EMs*. In particular, immune cells of the innate and acquired immune system play a key role in the survival and proliferation of endometrial cells outside the uterine cavity (4). However, its precise prevalence in the population is difficult to determine because it is asymptomatic or subclinical in most cases (5). Identifying molecular markers associated with *EMs* is of great significance for improving patient prognosis. Previous studies have suggested that several factors are involved in *EMs*, including genetic, and immune changes (1, 6). Previous researches indicated that Next-generation sequencing *(NGS)* may provide new clues in the pathogenesis of *EMs* (7). High-throughput *RNA*-sequencing *(RNA-Seq)* has provided new insight into the contribution of gene expression in disease contexts. Gene markers play an indispensable role in the prevention and diagnosis of diseases, offering novel perspectives for understanding the molecular mechanisms underlying these conditions and enabling the development of targeted therapeutics (8). In general, evaluating transcriptome datasets facilitates the assessment of overall gene functions and structures to investigate the molecular signatures predictive of certain diseases. Comparing the gene expression profiles of disease tissue to that of a normal healthy tissue is a powerful approach to understand the underlying cellular events in the etiology of any disease (9). Accordingly, previous studies have investigated differential gene expression through the use of diverse microarray platforms to explore molecular alterations associated with the *EMs* (10, 11). A total of 1309 upregulated and 663 downregulated genes were identified through the analysis of the transcriptomes of eutopic and ectopic endometrial stromal cells (12). The gene expression levels of *BCL6* and *LITAF* genes have been studied in the eutopic endometrial tissues of women with *EMs* in comparison with the normal endometrial samples (13). Significant expression differences were obtained for *SPARC, MYC, IGFBP1* and *MMP3* genes in the *Ems* (14). Identifying molecular diagnostic markers associated with *EMs* may allow for the early prediction of outcomes in patients and inform targeted treatment strategies, thereby substantially reducing the incidence of *EMs* (15). A large number of genes were identified in the occurrence of *EMs*, affecting immune system regulation, cell adhesion, and vascularization (2).

However, there are crucial challenges of individual research gene-level studies, such as heterogeneity among datasets, high cost, and relatively small dataset size, which could provide false positive or negative findings (16). Meta-analysis as a traditional method was implemented to successfully deal with the genetic study’s challenges, to identify reliable *EMs*-related biomarkers (17). Thus far, the exact reason for *EMs* is still not clear and, therefore meta-data analysis may provide further knowledge to solve the molecular pathogenesis complexity of such conditions. The genome wide association *(GWA)* meta-analysis was identified 5 novel loci significantly associated with *EMs* risk (18).

Furthermore, the biological processes that are involved with the differentially expressed genes *(DEGs)* and the functional enrichment analysis were also studied in *EMs* (19). Integrative-omics analysis identified critical roles of immune pathogenesis, Wnt signaling, differentiation, and migration of endometrial cells as hallmarks for *EMs* (19).

Gene co-expression network analysis has been used to extract new information using *DEGs* (20). Using weighted gene co-expression network analysis *(WGCNA),* Yang et al. constructed gene co-expression networks for multiple cancer types and found that some prognostic modules were conserved across different cancer types (21). During the *WGCNA* analysis, 18 co-expression modules were identified. Among them, the pale turquoise module contained the hub genes *FOSB, JUNB, ATF3, CXCL2*, and *FOS,* which showed a significant correlation with *EMs* (22).

Protein-protein interaction *(PPI)* networks have been widely used to characterize the underlying mechanisms of genes associated with complex diseases (23, 24). The majority of human diseases are caused by a group of correlated molecules or a network, rather than a single gene. Thus, identification and validation of biomarker networks is critical to disease diagnosis, prognosis and treatment (25).

In this study, we analyzed the expression profiles of *mRNA* using bulk *RNA* sequencing and predicted the related functions of up and down-regulated meta-*DE* genes by functional enrichment analysis. This is the first study to construct *mRNA* co-expression network by analyzing meta-*DE* genes in *EMs*. The results of functional enrichment analysis indicated that meta-DE gene were mainly enriched in *’Adherens junction,’ ‘p53 signaling pathway,’ and ‘AMPK signaling pathway.’* This study aims to identify key genetic factors and underlying molecular mechanisms involved in the pathogenesis of *EMs*. By leveraging bulk *RNA* sequencing data and integrative bioinformatics analyses, we successfully identified critical genes associated with *EMs*. The findings of this research not only enhance our understanding of the molecular basis of *EMs* but also contribute to the identification of novel key genes. Ultimately, early detection of *EMs* risk through molecular profiling can facilitate more treatment strategies and help minimize unnecessary clinical interventions.

## Materials and Methods

### Data collection

Bulk *RNA-Seq* datasets related to *EMs* were obtained from the Gene Expression Omnibus *(GEO)* database (https://www.ncbi.nlm.nih.gov/gds/). Three independent bulk *RNA-Seq* studies focusing on *EMs* were selected and analyzed. In the first dataset *(GSE130435)*, transcriptomic profiling of the endometrium was performed on samples collected from eleven women in the secretory phase of the menstrual cycle. Among them, six women were diagnosed with *EMs* (stage I-IV), and five control subjects had no evidence of the disease at the time of surgery for benign gynecologic disorders. The mean ages of the *EMs* and control groups were 37 and 42 years old, respectively (range 23–49 years). None of the patients had used hormonal therapy for at least three months prior to sample collection. Endometrial biopsies were obtained through the University of California San Francisco (UCSF) NIH Human Endometrial Tissue and DNA Bank, following approval by the *UCSF* Committee on Human Research *(IRB#10-02786)*, with written informed consent obtained from all participants. The primary aim of this analysis was to elucidate proinflammatory phenotypes of macrophages within the eutopic endometrium of women with *EMs*, with particular focus on a potential infectious contribution to disease etiology. *DEG* analysis was conducted between macrophages isolated from affected and unaffected women, revealing the involvement of distinct biological and signaling pathways. The altered macrophage phenotypes were associated with dysregulated gene expression in the eutopic endometrium, contributing to the establishment of a proinflammatory microenvironment. These changes are implicated in the pathogenesis of *EMs* and may underlie impaired reproductive outcomes observed in affected women. In the second dataset *(GSE134056)*, transcriptomic profiling of the endometrium was performed on samples obtained from 38 women aged between 18 and 49 years. Among them, 16 were diagnosed with *EMs* and 22 served as control. All participants underwent laparoscopic procedures, during which informed consent was obtained in accordance with Institutional Review Board (IRB) protocols. Endometrial biopsies were collected prior to surgery using suction pipelles (Cooper Surgical Uterine Explora Model I) under general anesthesia. Each biopsy yielded ≥250 mg of tissue, which was subsequently processed for high-throughput *mRNA* sequencing using the *Illumina NextSeq platform*. Samples were sourced from three different institutions: (1) Women’s and Children’s Hospital, University of Missouri; (2) Boone Hospital, Columbia, MO; and (3) University of California, San Francisco. To distinguish *EMs* cases from controls based on transcriptomic profiles, various supervised machine learning *(ML)* techniques were applied, including decision trees, partial least squares discriminant analysis *(PLS-DA),* support vector machines *(SVM),* and random forests. Furthermore, a generalized linear model *(GLM),* followed by a likelihood ratio test, was used to identify *DEGs* among the 14,154 genes analyzed. This analysis revealed 28 *DEGs*, of which 5 were upregulated and 23 were downregulated in *EMs* samples. Several of these genes were proposed as potential biomarkers for the disease. In the *GSE212787* study, two cohorts were analyzed to investigate the molecular mechanisms underlying *EMs,* with a particular focus on ubiquitination and its role in fibrosis development. All participants had normal menstrual cycles and had not received hormonal therapy in the three months prior to surgery. Cohort 1 included: Six control endometrial *(NC)* samples from non-endometriosis patients, six eutopic endometrial *(EU)* samples and ten ectopic endometrial *(EC)* samples from ten patients diagnosed with ovarian *EMs*. These samples underwent integrated transcriptomic and proteomic analyses using *RNA sequencing*. Samples were collected at the Department of Obstetrics and Gynecology, First Affiliated Hospital of Xiamen University, with ethical approval number KY2021-03. Cohort 2 consisted of: Five *NC* samples from non-*EMs* patients, Paired EU and EC endometrial samples from six patients with ovarian EMs. Label-free quantitative ubiquitylomics analysis was performed on these samples. They were collected from the Department of Obstetrics and Gynecology, First Affiliated Hospital of Fujian Medical University (Ethical approval: MRCTA, ECFAH of FMU [2021]484). For transcriptomic data analysis, *DEGs* was assessed using *DESeq2* with Benjamini– Hochberg false discovery rate *(FDR)* correction. Genes were considered significantly differentially expressed if they met the criteria of: Adjusted p-value *(FDR)* < 0.05 and Fold change *(FC)* > 2. To explore the biological significance of *DEGs*, Gene Ontology *(GO)* and *KEGG* pathway enrichment analyses were conducted. These analyses helped to identify key functional changes involved in the pathogenesis of *EMs*, particularly those associated with ubiquitination and fibrosis. Integrating these datasets through meta-analysis increases statistical power, reduces dataset-specific bias, and enables the identification of consistently dysregulated genes and pathways across diverse patient populations and experimental conditions. **Table 1** summarized the sample size, accession number, platform, Tissues, and references submitted to messenger *RNA-Seq*.

**Table 1.**
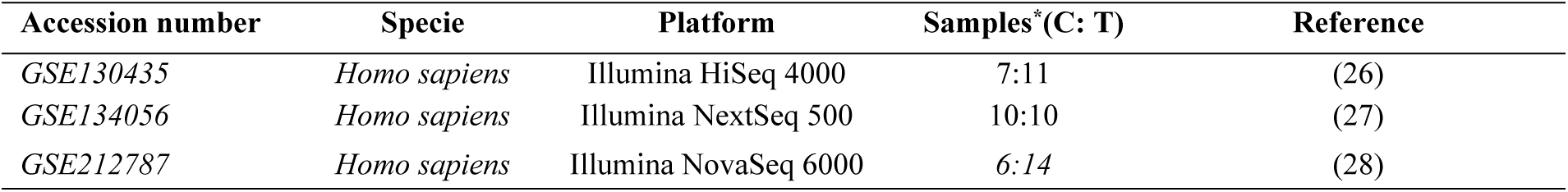
Bulk RNA-Seq datasets for transcriptomics analyses of endometriosis.

### Data processing and identification of *DE*Gs

Initially, read counts from patient and control groups were collected for transcriptomic analysis. Subsequently, *Ensembl IDs* within the count matrix were converted to *GeneIDs*. *DEGs* were identified using the *DESeq2* package (v1.28.1) implemented in *R* (29). *DESeq2* utilizes negative binomial generalized linear models to estimate gene-specific dispersion parameters. The statistical significance of *DEGs* was evaluated using the *Wald test* implemented within the *DESeq2* framework (30). Multiple testing correction was applied using the *FDR* method. Genes with a fold change ≥ |2| and an adjusted *P-value* ≤ 0.05 were considered significantly differentially expressed (31).

### Meta-Analysis of Datasets

Meta-analysis has been extensively utilized in genetic research, particularly for the identification of genes associated with various diseases (32). Subsequent to the identification of *DEGs*, a meta-analysis approach was employed to systematically integrate and synthesize data derived from multiple independent transcriptomic studies. This methodology enhanced the overall statistical power and analytical robustness, thereby facilitating the identification of meta-genes that demonstrate significant associations with the *EMs*. To perform the meta-analysis, two widely *P-value* combination methods: *Fisher’s* method and the *inverse* normal method were implemented using the *metaRNASeq* package (v1.0.5) from the Bioconductor (33).

### Biological Pathway Enrichment Analysis

Pathway enrichment analysis helps researchers gain mechanistic insight into gene lists generated from omics experiments. This method identifies specific biological pathways *(BPs),* molecular functions *(MFs),* cellular component *(CCs)* and Kyoto Encyclopedia of Genes and Genomes pathways *(KEGG)* that are enriched in a group of *DEGs* (34, 35). For enrichment analysis, the list of meta-genes was imported to *Enrichr* database. Detecting over-represented biological and molecular pathways donates valued comprehension of biological mechanisms in *EMs*.

### Construction of gene co-expression networks

The *WGCNA* Bioconductor R package (v3.5.1) was employed to construct gene co-expression networks, detect the correlation patterns among genes and identify important modules across bulk *RNA-Seq* samples (36). The expression values of meta-genes were normalized using the variance-stabilizing transformation function *(vst)*, in order to generate a matrix of values for which variance is constant across the range of mean values (37). The *stringsAsFactors* function was used while checking for missing values in the dataset. Hierarchical clustering using the *hclust* function was applied to detect outlier samples. The scale-free topology features of biological networks were incorporated through the application of the *pickSoftThreshold* function. The adjacency matrix was constructed by employing Pearson correlations methods across meta-genes (20, 38). Afterwards, the adjacency matrix was converted into a Topological Overlap Matrix *(TOM)* and corresponding dissimilarity matrix *(1−TOM)* for the identification of gene modules for each pair of genes with strong interconnectivity (36). Finally, the *cutreeDynamic* function, along with average linkage hierarchical clustering, was used to clustered genes into modules based on similar expression patterns. To determine the effect of hub genes on the modules, they were imported to *Cytoscape*.

### Network Topological Analysis

Topological data analysis represents a systems biology-oriented approach that has emerged as one of the most powerful methodologies for the investigation and interpretation of complex biological networks (39). The topological analysis of Protein-protein interactions *(PPIs)* network assist to explore the molecular mechanisms and pathways regulated by the essential genes in an organism (40). *PPI* among meta-genes will be analyzed and visualized using *Cytoscape* software to construct gene interaction networks. Hub genes will be identified based on their degree of connectivity and their relative biological significance compared to other genes within the network (41). In the network the nodes correspond to proteins and the edges to the interactions between each protein. We then used the *CentiScaPe Cytoscape* plug-in to calculate the node degree and betweenness centrality of each protein. The nodes (proteins) that had a degree centrality and a betweenness centrality greater than or equal to the mean were identified as key proteins more likely to modulate symptoms of *EMs*.

### Prediction of *miRNAs* targeting the hub genes

Computational prediction of *miRNAs* plays a pivotal role in the early stages of research, as it provides essential insights into the complex regulatory networks governed by these small noncoding *RNAs*. Candidate *miRNAs* targeting the hub genes were systematically predicted using the *miRDB* database.

## Results

### Identification of *DEGs*

The step-by-step workflow of the systems biology approach used in this study is presented in **Fig 1**. Initially, a comprehensive analysis was conducted on a total of 58 endometrial tissue samples, from three independent bulk RNA-Seq datasets. These samples included 35 cases diagnosed with *EMs* and 23 control samples from healthy individuals.

**Fig 1.**
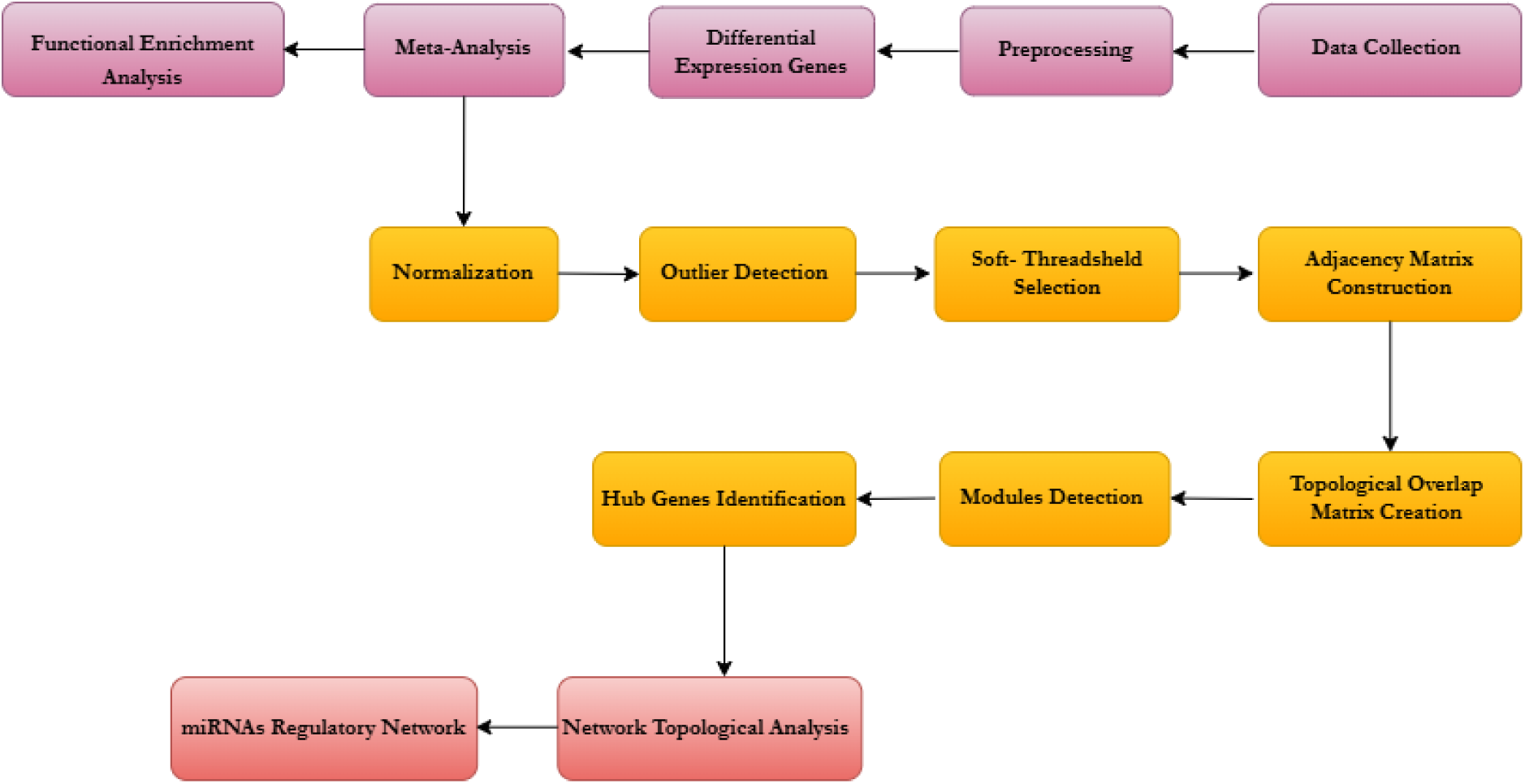
Workflow of the systems biology methodology in this project

After pre-processing of the bulk *RNA*-seq data of each studies, we created datasets containing the genes of 1117, 1005, 3123 in *GSE130435, GSE134056*, and *GSE212787*, respectively **(Fig 2)**. We performed differential analysis using the *GLM* followed by likelihood ratio test on 5245 genes and found 2005 upregulated and 3240 downregulated genes overall.

**Fig 2.**
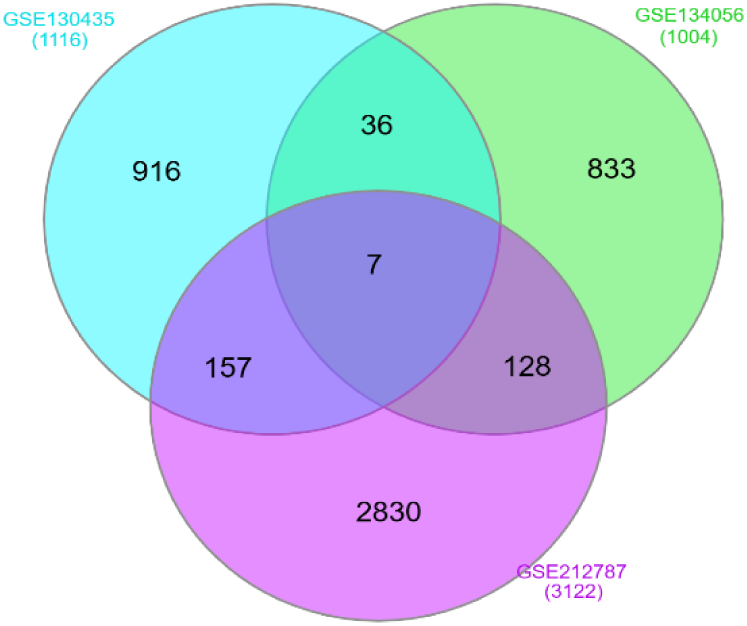
Identification of DEGs Across Three Datasets via Venn Diagram Analysis

### Meta-analysis of *DEGs*

Overall, 603 and 443 meta-genes, were discovered in response to *EMs* using the *Fisher* and *Invorm* methods, respectively. Among these, 427 common meta-genes considered for identifying correlation patterns. **Fig 3** displays the outcomes of meta-analysis of bulk *RNA-Seq* datasets.

**Fig 3.**
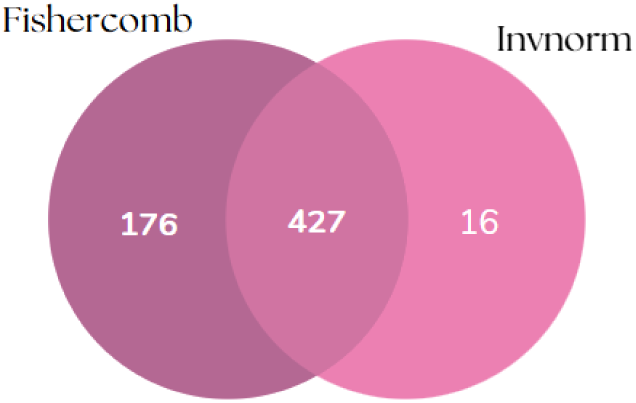
Identification of 427 Meta-Genes Using Fisher and Invorm

### Functional Enrichment Analysis of meta-genes

Functional enrichment analysis was performed on the identified meta-genes to elucidate associated *BPs, MFs, CCs,* and signaling pathways. The results of this comprehensive analysis are illustrated in **Fig 4** and **Table 2**, highlighting the key functional categories and enriched pathways potentially implicated in the underlying biological mechanisms.

**Fig 4:**
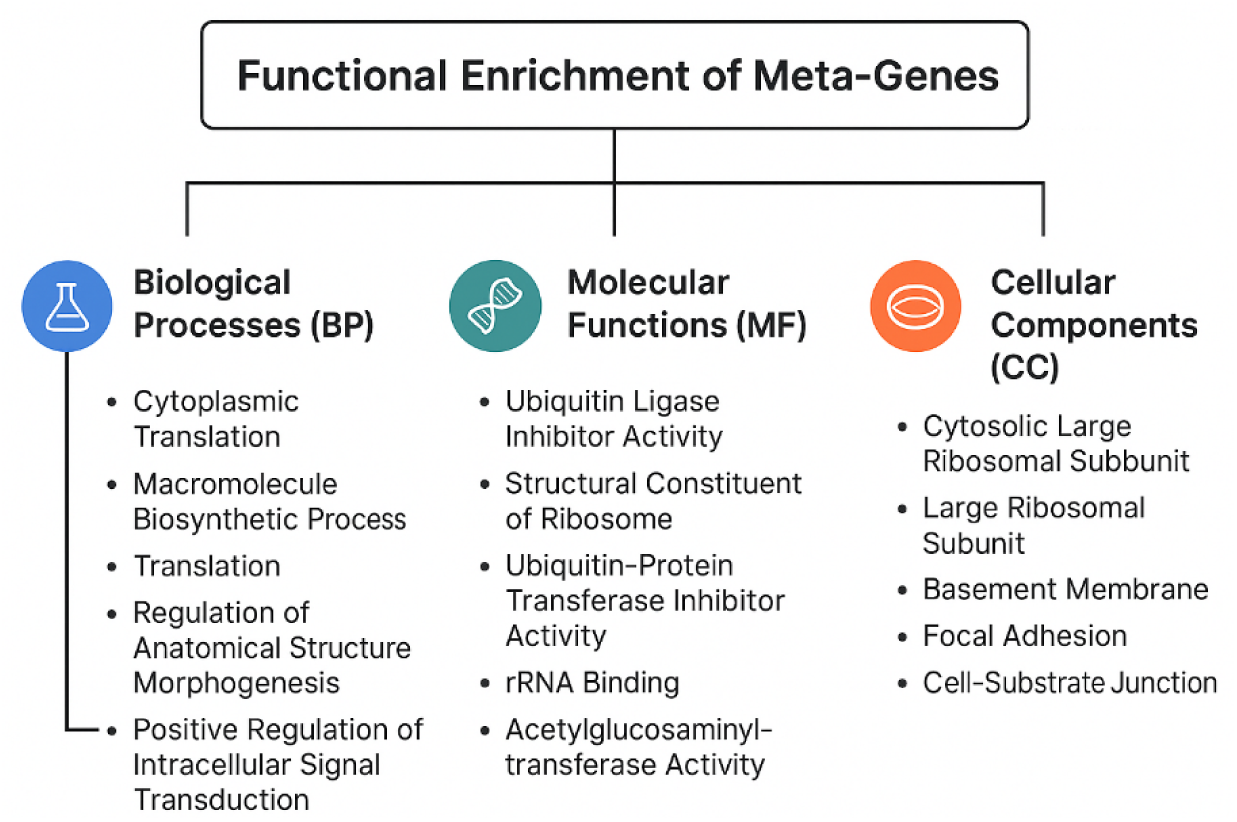
Functional Enrichment Analysis of Meta-Genes across *BPs, MFs,* and *CCs*

**Table 2:** The most significant pathways identified through the analysis of meta-genes in EMs.

| Pathway name | P-value | Total genes in pathway |
| --- | --- | --- |
| Pyruvate metabolism | 0.003131 | 6 |
| Central carbon metabolism in cancer | 0.005840 | 7 |
| Thyroid hormone signaling pathway | 0.012946 | 9 |
| Human papillomavirus infection | 0.015325 | 18 |
| Propanoate metabolism | 0.020199 | 4 |
| Adherens junction | 0.022501 | 6 |
| p53 signaling pathway | 0.025414 | 6 |
| Protein export | 0.032977 | 3 |
| AMPK signaling pathway | 0.033036 | 8 |
| Glucagon signaling pathway | 0.048457 | 7 |

### Construction of the gene co-expression network

In order to comprehensively investigate the functional relationships and co-regulatory patterns among the identified meta-genes associated with *EMs*, we employed *WGCNA* software. A crucial step in the construction of network topology is the selection of an appropriate soft-thresholding power (β), which influences the strength of correlation between gene pairs. We selected β = 20 as the optimal power, since it was the lowest value at which the scale free topology fit index (R²) reached 0.9, indicating a strong scale-free topology **(Fig 5)** (36).

**Fig 5:**
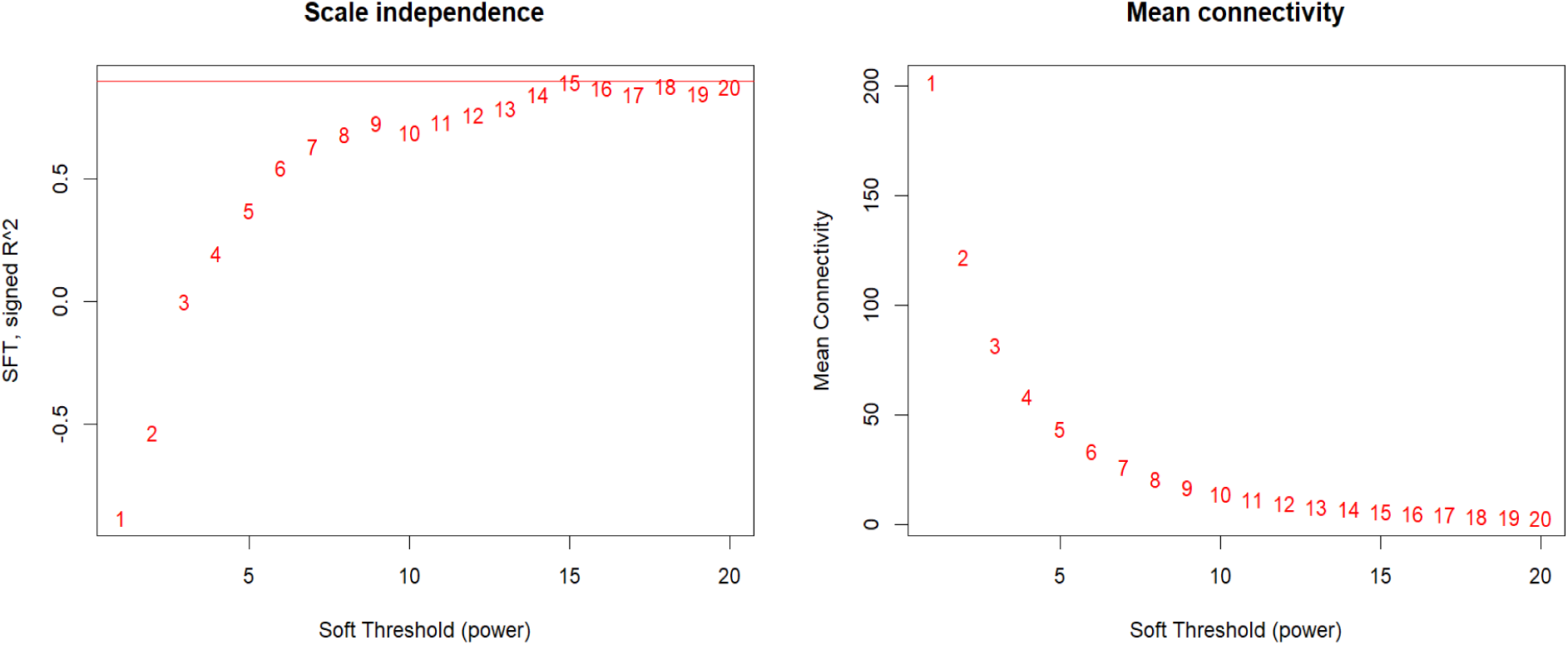
Scale independence and mean connectivity of network topology for different soft-thresholding

Meta-genes with similar expression patterns were clustered into co-expression modules that were displayed in different colors. A total of four distinct gene co-expression modules were identified. The largest module in the network is significant and contains 205 genes, while the non-significant modules consist of 170, 41, and 11 genes **(Fig 6A).** The relevance between each gene co-expression module and clinical information was further explored through module–trait relationship analysis. The resulting visualizes the strength and direction of these correlations, with each cell representing the Pearson correlation coefficient and the associated *P-value* **(Fig. 6B).** Among these, the turquoise module, containing 205 genes, was found to have a significant correlation with *EMs* (R = 0.36, P = 0.005). This module was considered for downstream analyses, including the identification of hub genes and the exploration of potential regulatory mechanisms involved in *EM*-related pathways. To explore the biological relevance of modules, the module–trait relationships were assessed by correlating the module eigengenes with the phenotypic trait ‘weight’. Additionally, gene significance *(GS)* for weight was calculated and integrated into the visualization, allowing the identification of modules most strongly associated with the trait **(Fig 6C).** To visualize the topological structure of the gene co-expression network, a network heatmap plot was constructed based on the *TOM.* Prominent blocks of darker coloration along the diagonal were observed, indicating regions of high interconnectivity that correspond to modules identified through hierarchical clustering. These coherent clusters reflect elevated intramodular connectivity and suggest that genes within each module may be functionally related or co-regulated **(Fig. 6D).**

**Fig 6:**
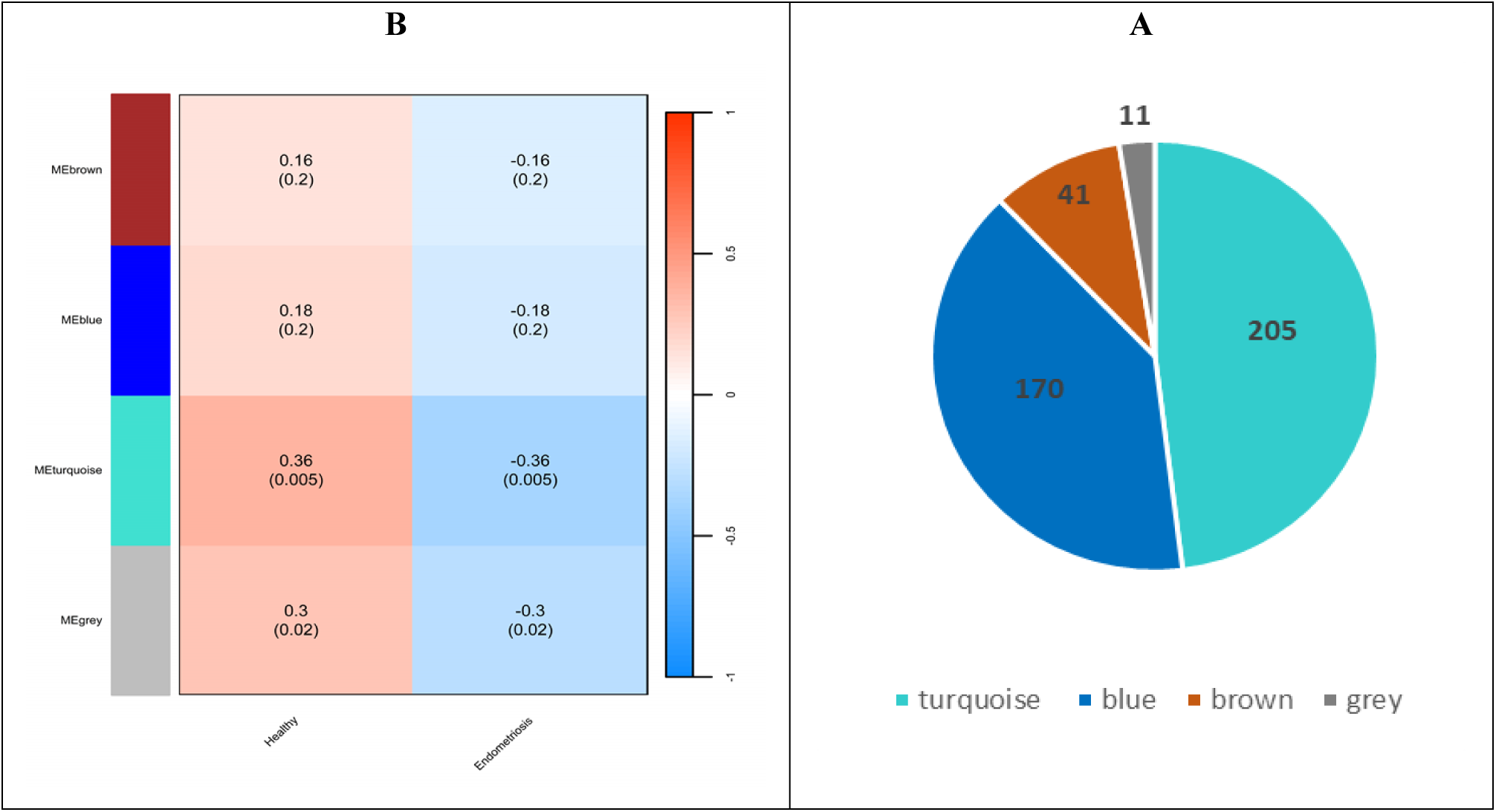
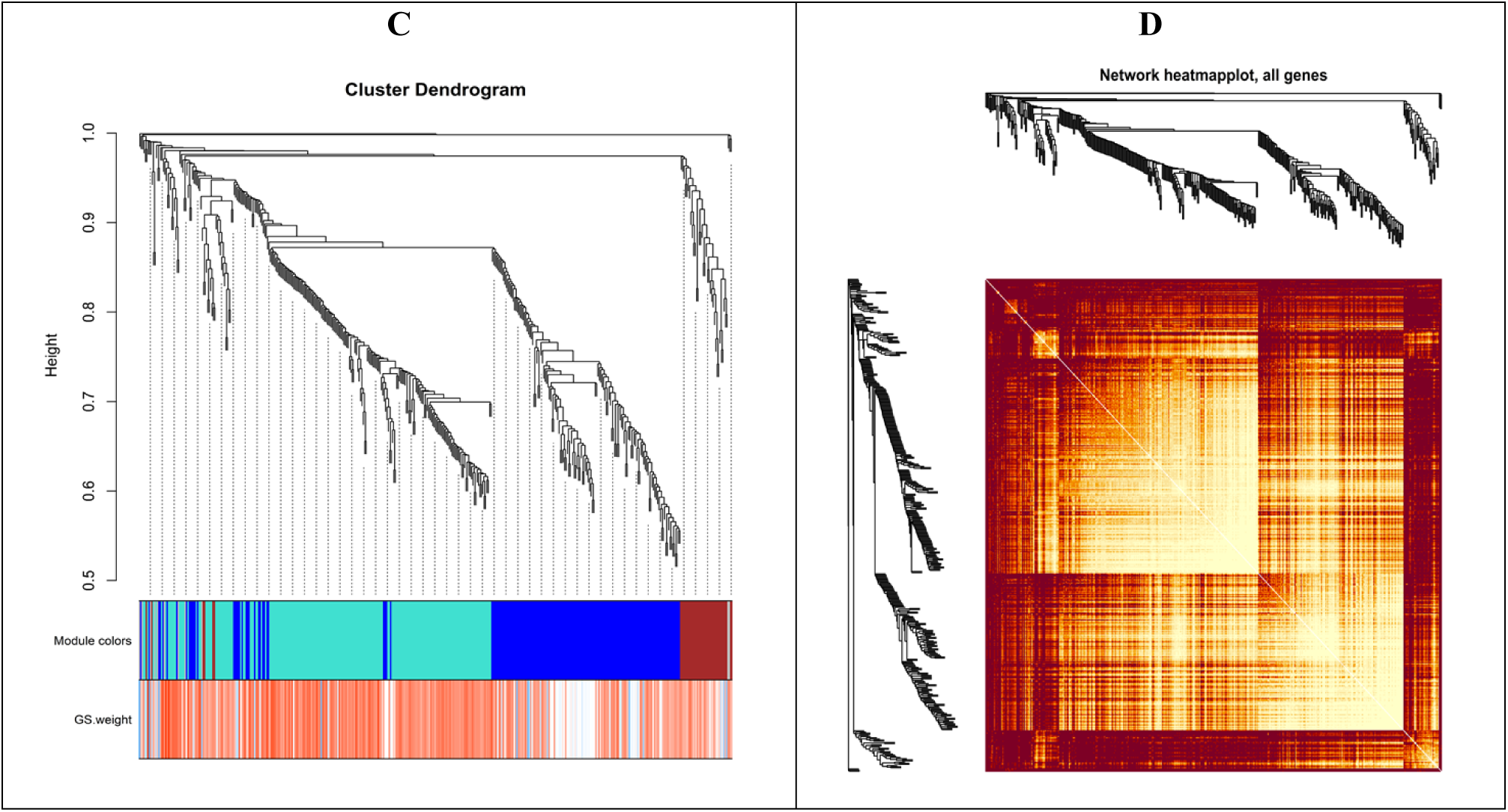
A) The sizes of determined modules based on the number of involved genes. B) Determination of module-trait relationship of EMs and identification of the most clinically relevant modules; each row indicates a module eigengene, and each column represents a clinical trait. C) Cluster dendrogram of the meta-genes. The branches and color bands demonstrate the specific module. D) TOM plot; light color symbolizes low overlap, and progressively darker red color symbolizes higher overlap between common genes. Blocks of darker colors along the diagonal correspond to modules.

Using the density-based clustering non-parametric algorithm *(DBSCAN),* we analyzed the 20 hub genes identified within the turquoise module of co-expression network. These genes included *NKAP, ZFTA, OGN, CEP112, TEF, JCAD, IGFBP3, SCD5, OLFML1, CC2D2A, XYLT2, ME3, ANK2, KRBA1, NLGN3, PLCD3, LRRC17, PRKG1, ZFP2,* and *PTPRB*. The *STRING* database was employed to assess network among these hub genes.

### PPI of the significant module

Using the *Cytoscape* platform, a comprehensive *PPI* network was constructed based on the 205 *DE*-meta genes identified. The value of the Betweenness Centrality *(BC)* is between 0 and 1. The node size in the networks represents the centrality of the corresponding nodes. As shown in **Fig 7**, the *PPI* network was observed to represent the central hub positions of several genes. Each node was designated to represent a protein, and each edge between them was used to indicate an interaction between two proteins. Smaller and lighter-colored nodes correspond to proteins with fewer interactions, which may perform more specific and limited roles in biological processes. Collectively, these genes are expected to contribute to key biological processes including signal transduction, transcriptional regulation, cellular metabolism, and structural remodeling. Proteins such as *RPAP2, TBX1, LRRC49, MAS1, TTTL7, MSI2, SERPINE, TONSL, STAR, SVIL, PLCD3, LIPT1, TPH1*and *PIPOX* were commonly identified as hub proteins due to their extensive interactions with numerous other proteins within the network. These proteins, which were characterized by a higher degree of interconnectedness and putative co-involvement in discrete biological pathways, were systematically partitioned into functionally relevant clusters, as delineated through integrative genetic network analyses. These hub proteins were found to play crucial roles in maintaining network integrity and regulating essential biological processes. Hub proteins have been proposed as critical regulators in endometriosis, highlighting their potential functional significance in the underlying biological processes. For example, *MSI2* has been implicated in *RNA* binding and stem cell maintenance (42), whereas *PLCD3* is involved in phosphoinositide signaling a pathway frequently related to cell proliferation and migration (43). Additionally, the presence of transcriptional regulators such as *RPAP2* and chromatin-modifying enzymes like *KAT14* suggests potential epigenetic modulation within the disease context (44). The identification of *STAR* and *MAS1*, are known to participate in steroidogenesis and hormonal signaling, further indicates the possible involvement of endocrine regulatory mechanisms (45).

**Fig 7.**
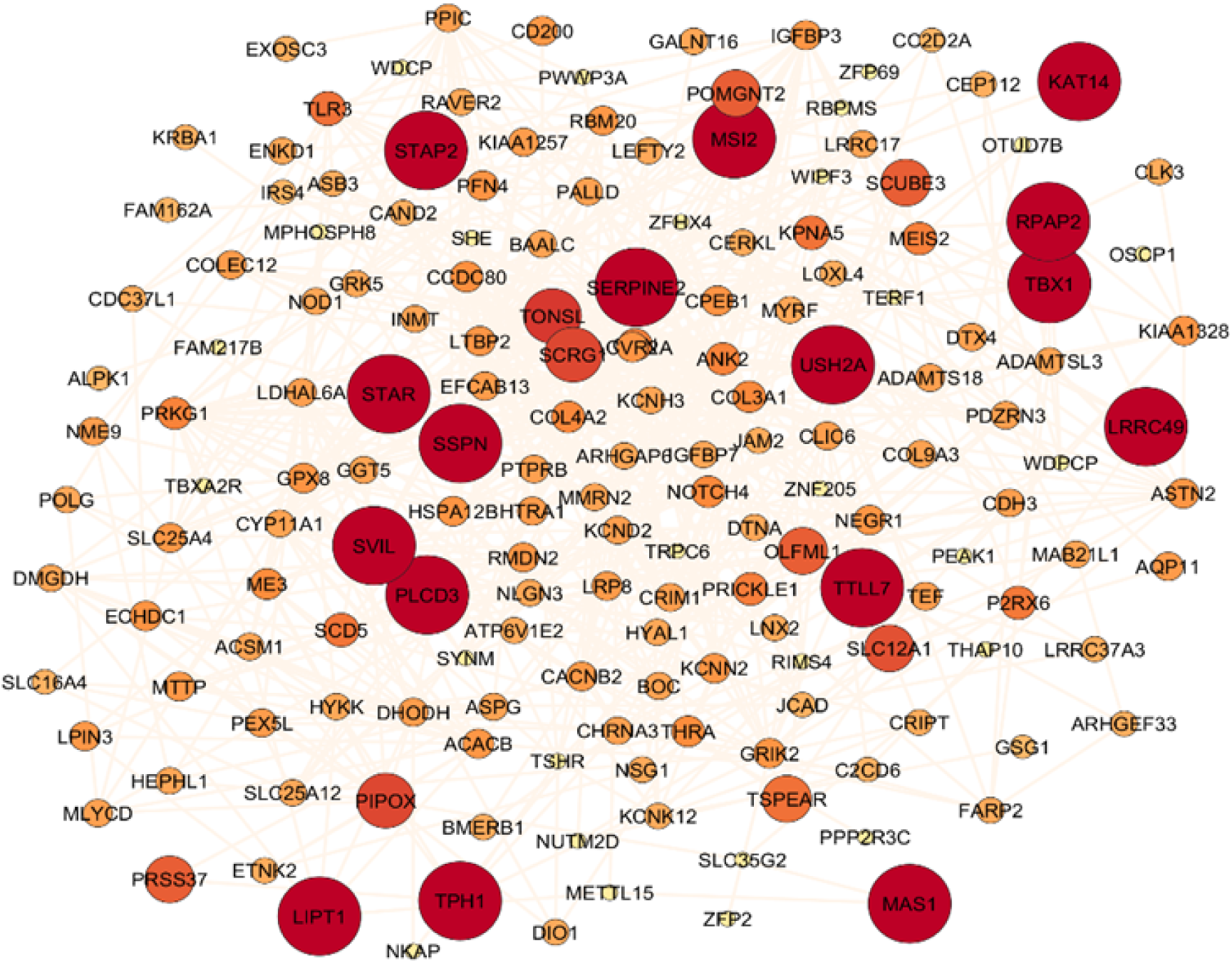
Protein–protein interaction network for the meta-genes using *Cytoscape*.

### Construction of the *miRNAs* Regulatory Network

Using the *miRDB* database, we identified 39, 53, 9, 516, 35, 43, and 101 *miRNAs* targeting *TBX1, RPAP2, MAS1, MSI2, STAR, PLCD3,* and *SVIL,* respectively. These findings suggest that the identified *miRNAs* may regulate distinct regulatory networks by targeting hub genes, thereby modulating critical signaling pathways and biological processes involved in the progression of *EMs*. The hub gene *MSI2* was found to share twenty-nine common *miRNAs* with *SVIL*, as illustrated in **Fig 8**, highlighting a substantial overlap in their post-transcriptional regulatory networks and suggesting that these two genes may be co-regulated by a similar set of *miRNA* molecules.

**Fig. 8.**
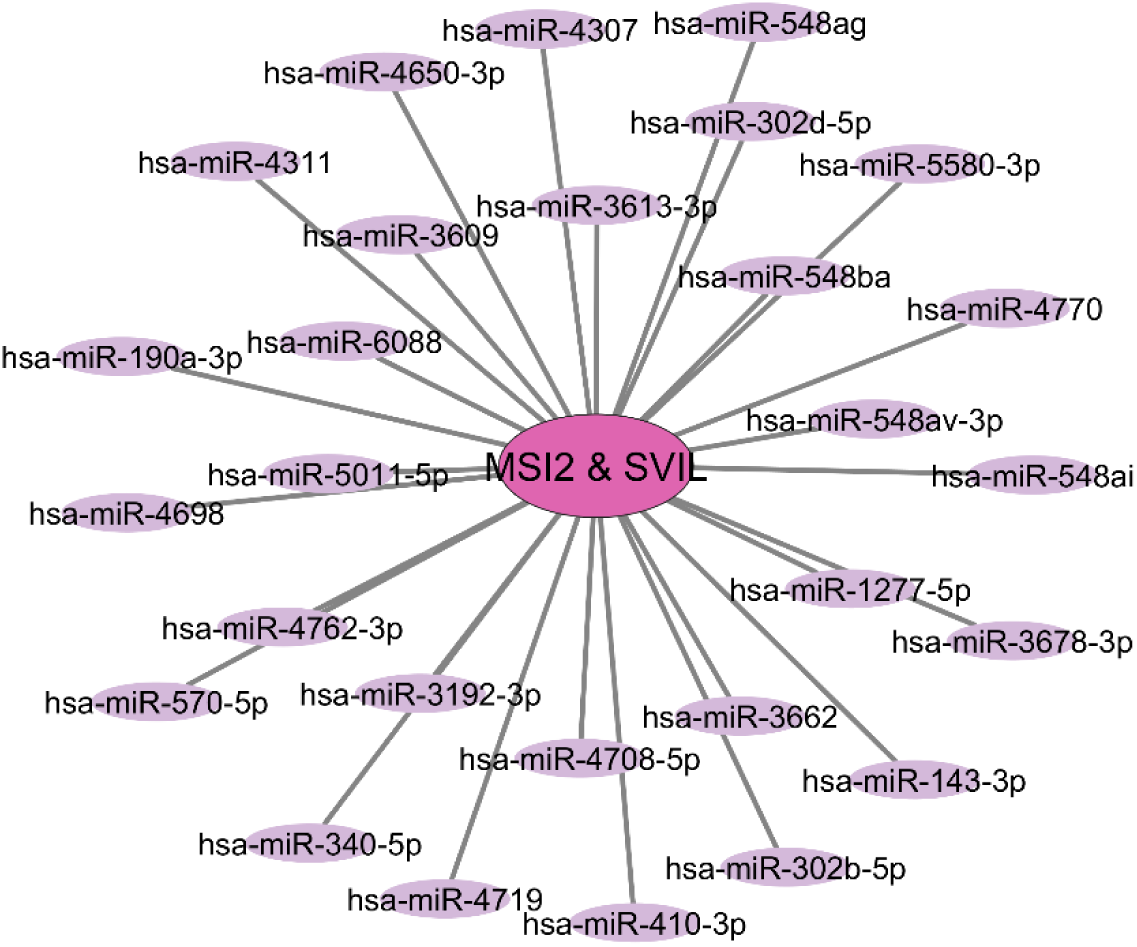
Network visualization of *MSI2 & SVIL* and their common associated *miRNAs*, constructed in *Cytoscape*.

## Discussion

The availability of bulk *RNA-seq* has enabled a more comprehensive characterization of molecular alterations underlying various diseases. *EMs* remains a significant challenge in reproductive medicine. This study aims to identify key hub genes involved in *EMs* using a bulk *RNA-seq* analysis approach. Initially, relevant datasets were obtained from the *GEO* database. After normalization, *DEGs* associated with *EMs* were identified. To achieve sufficient statistical power and gain novel insights into the relationships and expression patterns of key regulatory genes, a meta-analysis was conducted. This approach aimed to identify meta-genes consistently dysregulated across multiple datasets, which may serve as potential biomarkers. To further explore the functional implications of these genes, pathway and *GO* enrichment analyses were performed, aiming to uncover the critical biological functions and pathways involved in *EMs*. Subsequently, we employed *WGCNA* to construct gene co-expression networks and identify gene modules associated with the disease. In addition, a disease-specific interaction network for *EMs* was constructed. This comprehensive approach allowed us to pinpoint hub genes and potential biomarkers relevant to the diagnosis and prognosis of *EMs*. Unlike previous studies that primarily relied on microarray-based datasets, our research integrates bulk *RNA-seq* data with differential expression analysis, meta-analysis, *WGCNA*, and *PPI* network analysis. Through this integrative approach, we identified key genes under this specific conditions that have not been previously reported, offering novel targets for early diagnosis and therapeutic intervention in *EMs*.

Overall, the meta-analysis approach detected 427 common meta-genes using the *Fisher* and *Invorm* methods, which missense mutations in the *PTEN* meta-gene have been reported to contribute to the molecular mechanisms underlying endometriosis, as well as the genotypic-phenotypic correlations observed in endometrial and ovarian cancers (46). Serial analysis of gene expression revealed differential expression of the *B3GNT5* meta-gene between endometriosis and normal endometrium (47).

Our enrichment analysis revealed that these meta-genes are predominantly associated with several key biological pathways and disease processes. The involvement of these pathways suggests potential mechanistic associations with *EMs*, particularly in relation to altered cellular metabolism, and immune dysregulation. Analysis of exosomal *microRNAs* revealed that anatomical structure morphogenesis is a significantly enriched biological process associated with the pathogenesis of *EMs* (48). The process of positive regulation of intracellular signal transduction significantly influences the molecular mechanisms underlying the development of *EMs* (49). Recent research highlighted the essential association between ubiquitination and *EMs* pathogenesis, (50), also, Acetylglucosaminyltransferase Activity is involved in glycosylation, which affects cell signaling, adhesion, and immune interaction, all processes relevant to *Ems* (51). The association of the *p53 signaling pathway* with *EMs* susceptibility has been investigated in a Taiwanese population (52). In *EMs*, dysregulation of the *p53 pathway* may contribute to enhanced cell survival, resistance to apoptosis, and the invasion of endometrial-like tissue beyond the uterine cavity (53). The *AMPK signaling pathway* plays a crucial role in regulating cellular metabolism, inflammation, and the immune response, the dysregulation of *AMPK signaling* has been implicated in several female reproductive disorders, including *EMs*, infertility, and reproductive ageing (54). Targeting *EphA2* has been reported to inhibit the progression of *EMs* by modulating the *AMPK signaling pathway* (55). *Adherens junctions* have been reported to decline during the preimplantation period, potentially facilitating trophoblast invasion through the epithelial barrier (56). Dysfunction of the *Adherens junction* appears to contribute to the detachment of endometriotic cells, representing a key initial step in the development of ovarian *EMs* (57).

In the current study, systems biology analysis revealed distinct topological characteristics within the co-expression network of *EMs*-related genes. Notably, *IGFBP7, IGFBP3,* and *NKAP* were grouped within a turquoise co-expression module, which was selected for discussion due to its potential relevance to endometriosis pathophysiology. The multifaceted functions of *IGFBP7,* including its regulatory roles in cell proliferation, apoptosis, and migration, were investigated to elucidate the underlying mechanistic pathways. Notably, *IGFBP7* appears to play a crucial role in the pathogenesis of endometriosis (58, 59). *MicroRNA-210-3p* has been identified as a critical post-transcriptional regulator in endometriosis, contributing to the development and progression of endometriotic lesions by directly targeting *IGFBP3.* Through the downregulation of *IGFBP3, miR-210-3p* may enhance cellular proliferation, migration, and survival within ectopic endometrial tissue, thereby promoting lesion establishment and maintenance (60). *NF-κB*, also known as *NKAP,* has been shown to be activated in peritoneal endometriosis in women, underscoring its pivotal role in promoting inflammation within ectopic lesions. This activation supports the notion of a distinct inflammatory microenvironment in endometriotic implants compared to surrounding normal tissue. The *NF-κB* signaling pathway is believed to contribute to the persistence and progression of endometriotic lesions by regulating pro-inflammatory cytokines, immune cell recruitment, and the expression of adhesion and survival-related genes (61). This study highlighted several genes including *ZFTA, OGN, CEP112, TEF, JCAD, SCD5, OLFML1, CC2D2A, XYLT2, ME3, ANK2, KRBA1, NLGN3, PLCD3, LRRC17, PRKG1, ZFP2,* and *PTPRB* that may represent novel molecular contributors or potential biomarkers for endometriosis. However, further research involving experimental validation and detailed functional analyses is necessary to clarify their precise roles in the disease’s pathophysiology.

The gene network visualization of the *DE*-meta gene signatures are represented in **Fig 7**. The significance level for the hub genes is set at BC ≥ 0.1. *TBX1* was identified as an upregulated transcription factor expressed differential expression in *EMs*, (62), suggesting its potential involvement in the underlying molecular mechanisms of reproductive dysfunction. *TBX1* is a transcription factor involved in tissue development and immune system regulation (63). Alterations in this gene may be observed in tissue abnormalities and endometriosis. *RPAP2* has been implicated in the regulation of gene transcription, and its alteration may be associated with the dysregulation of genes involved in inflammation and tissue growth, (64) which can be considered relevant to endometriosis. It has been demonstrated that *MAS1* is expressed in the eutopic proliferative endometrium of patients with ovarian endometriotic tissues. This suggests that *MAS1* may be involved in the initiation of endometriosis, particularly in the migration of endometrial tissues from eutopic to ectopic sites (65). It was also observed that *MSI2* is downregulated as a result of the overexpression of *miR-145*, a molecule known to be dysregulated in endometriosis. This downregulation leads to alterations in cell behavior, including increased invasiveness, as demonstrated by the Matrigel Invasion Assay (66). The results demonstrate that aberrant expression of *STAR* in ectopic endometriotic tissues, resulting in increased peritoneal progesterone levels, is associated with the development of endometriosis (45). Current studies suggest that the endometriosis susceptibility locus on *17q21* may be associated with *PLCD3* mapping and the involvement of *PI-PLC δ3* in the disease (67). Gene expression analysis indicated that *LIPT1* expression was downregulated in uterine corpus endometrial carcinoma *(UCEC)* and was identified as a cuproptosis-related prognostic marker (68). The observed expression patterns suggest that *SVIL* is consistently expressed a strong to moderate levels in both normal endometrial tissues and endometrial cancer *(EC)* tissues (69).

*hsa-miR-143-3p* has been frequently reported as dysregulated in ectopic and eutopic endometrial tissues, and appears to contribute to the abnormal cell proliferation, migration, and invasion characteristic of *Ems* (70). Likewise, *hsa-miR-340-5p* shows altered expression in endometriotic lesions, and is supposed to participate in inflammatory and proliferative signaling pathways associated with disease progression (71). *hsa-miR-410-3p* is commonly downregulated in *EMs* and is thought to influence key pathways involved in cellular proliferation and immune responses (72). In addition, *hsa-miR-302b-5p*, implicated in endometrial stem cell regulation, has emerging evidence linking it to the pathophysiology of *EMs* (73).

## Conclusion

Through integrative systems biology analysis, our study successfully identified hub genes that may serve as potential diagnostic biomarkers for *EMs*. These findings provide novel insights into the genetic and molecular mechanisms underlying *EMs*, helping to fill a critical gap in current research. By characterizing gene expression profiles and associated signaling pathways, our results establish a theoretical framework for early disease prediction and the development of personalized therapeutic strategies. However, several limitations must be acknowledged. First, the diagnostic potential and biological significance of the identified hub genes have not yet been validated through experimental studies or machine learning-based predictive modeling. Additionally, the specific molecular functions and regulatory roles of these genes in the pathogenesis and progression of *EMs* remain unclear and require further investigation. Moreover, as our conclusions are based on bioinformatics analysis, validation through in vitro, in vivo, or clinical studies is essential. Despite these limitations, the identification of robust genetic markers in this study holds promise for improving early diagnosis, optimizing treatment plans, and potentially reducing the incidence of *EMs* in the future.

## Acknowledgments

We sincerely thank Dr. Mohammad Farhadian for his valuable assistance in our project.

